# Architecture of fully occupied Glua2 Ampa receptor – TARP complex elucidated by single particle cryo-electron microscopy

**DOI:** 10.1101/060046

**Authors:** Yan Zhao, Shanshuang Chen, Craig Yoshioka, Isabelle Baconguis, Eric Gouaux

**Affiliations:** Vollum Institute, Oregon Health & Science University, Portland, Oregon 97239, USA; Department of Biomedical Engineering, Oregon Health and Science University, 2730 SW Moody Ave, Portland OR 97201; Howard Hughes Medical Institute, Oregon Health & Science University, Portland, Oregon 97239, USA

## Abstract

Fast excitatory neurotransmission in the mammalian central nervous system is largely carried out by AMPA-sensitive ionotropic glutamate receptors. Localized within the postsynaptic density of glutamatergic spines, AMPA receptors are composed of heterotetrameric receptor assemblies associated with auxiliary subunits, the most common of which are transmembrane AMPA-receptor regulatory proteins (TARPs). The association of TARPs with AMPA receptors modulates the kinetics of receptor gating and pharmacology, as well as trafficking. Here we report the cryo-EM structure of the homomeric GluA2 AMPA receptor saturated with TARP γ2 subunits, showing how the TARPs are arranged with four-fold symmetry around the ion channel domain, making extensive interactions with the M1, M2 and M4 TM helices. Poised like partially opened ‘hands’ underneath the two-fold symmetric ligand binding domain (LBD) ‘clamshells’, one pair of TARPs are juxtaposed near the LBD dimer interface, while the other pair are near the LBD dimer-dimer interface. The extracellular ‘domains’ of TARP are positioned to not only modulate LBD ‘clamshell’ closure, but also to affect conformational rearrangements of the LBD layer associated with receptor activation and desensitization, while the TARP transmembrane (TM) domains buttress the ion channel pore.

---

Fast excitatory neurotransmission at chemical synapses of the brain underpins a spectrum of activities ranging from memory and learning, to speech and hearing, to movement and coordination. Ionotropic glutamate receptors (iGluRs) are a family of transmitter-gated ion channels comprised of three related subfamilies – AMPA, kainate and NMDA receptors – that mediate the majority of ionotropic excitatory signaling^1^. Neuronal AMPA and kainate receptors, by contrast with NMDA receptors, are associated with auxiliary membrane protein subunits that, in turn, modulate receptor gating, trafficking, and pharmacology^2^.

Stargazin is the founding member of the transmembrane AMPA receptor regulatory proteins (TARP)^3^, a family of membrane proteins related in amino acid sequence to claudin, a four-helix transmembrane protein^4^. Coexpression of recombinant AMPA receptors with TARPs largely recapitulates native receptor gating kinetics, ion channel properties, and pharmacology, consistent with the notion that TARPs are fundamental components of neuronal AMPA receptor signaling complexes^5^, yet with a heterogeneous stoichiometry ranging from 1 to 4 TARPs per receptor^6^. Stargazin, also known as TARP γ2, modulates AMPA receptor gating by slowing deactivation and desensitization, accelerating the recovery from desensitization, increasing the efficacy of partial agonists such as kainate, and attenuating polyamine block of calcium-permeable AMPA receptors^7,8^. Despite progress in visualization of the AMPA receptor – TARP complex at a low resolution^9^, determination of the molecular architecture of the AMPA receptor – TARP complex and defining a molecular mechanism for TARP modulation of receptor function have proven elusive, in part because TARPs are bound weakly to the receptor and dissociate under typical conditions employed in complex solubilization and purification.

X-ray crystal and single particle cryo-electron microscopy (cryo-EM) structures of AMPA receptors show that they are tetrameric assemblies consisting of three layers – the amino-terminal domain (ATD), the ligand-binding domain (LBD) and the trans-membrane domain (TMD)^10–14^. Whereas the ATDs and LBDs assemble as two-fold symmetric dimers-of-dimers^15,16^, the TMDs adopt four-fold symmetry, thus resulting in a symmetry mismatch between the TMD and the LBD and giving rise to two-fold related, conformationally distinct subunit pairs, A/C and B/D^10^. Each LBD resembles a clam-shell^17^, that is open in apo and antagonist-bound states and closes upon binding of agonists^18^. Structures of the GluA2 receptor in agonist-bound, pre-open states shows that the LBDs are assembled in a ‘back-to-back’ fashion, with agonist-induced closure of the LBDs causing a separation of the LBD-TMD linkers and a translation of the LBD layer closer to the membrane^11,12^. The agonist-bound desensitized state, by contrast, undergoes a massive rearrangement of the ATD and LBD layers, thus decoupling agonist-binding from ion channel gating^12,13,19^.

To define the molecular basis for TARP modulation of AMPA receptor gating and pharmacology, we sought to elucidate the architecture of the AMPA – TARP complex by single particle cryo-EM. Here we focus on the wild-type, homomeric rat GluA2 AMPA receptor^20^, bearing an arginine at the Q/R site^21^ and harboring the flop splice variant^22^, where we have coexpressed the receptor in mammalian cells in combination with full-length TARP γ2^23^. Evidence for formation of a physiologically relevant receptor-TARP complex in these cells was shown by a diagnostic increase in the efficacy of the partial agonist, kainate, to 80±2% of that of a full agonist, glutamate^24^ (Fig. 1a). To define conditions for solubilization and purification of AMPA receptor fully bound with TARPs, we carried out fluorescence-detection size-exclusion chromatography (FSEC)^25^ studies on mammalian cells co-expressing GluA2 receptor and an engineered TARP γ2-eGFP fusion^26^. By systematic screening of detergents and lipids via FSEC, we found that whereas dodecyl maltopyranoside (DDM) leads to dissociation of the receptor – TARP complex, digitonin retains the complex integrity, allowing TARP to remain associated with receptor following solubilization and purification (Extended Data Fig. 1a). We proceeded to purify the native GluA2 receptor-full length TARP complex in the presence of the competitive antagonist MPQX^27^ (Extended Data Fig. 1b and 1c), succeeding in isolating a homogeneous population suitable for single particle cryo-EM analysis (Extended Data Fig. 1d and 1e).

**Figure 1.**
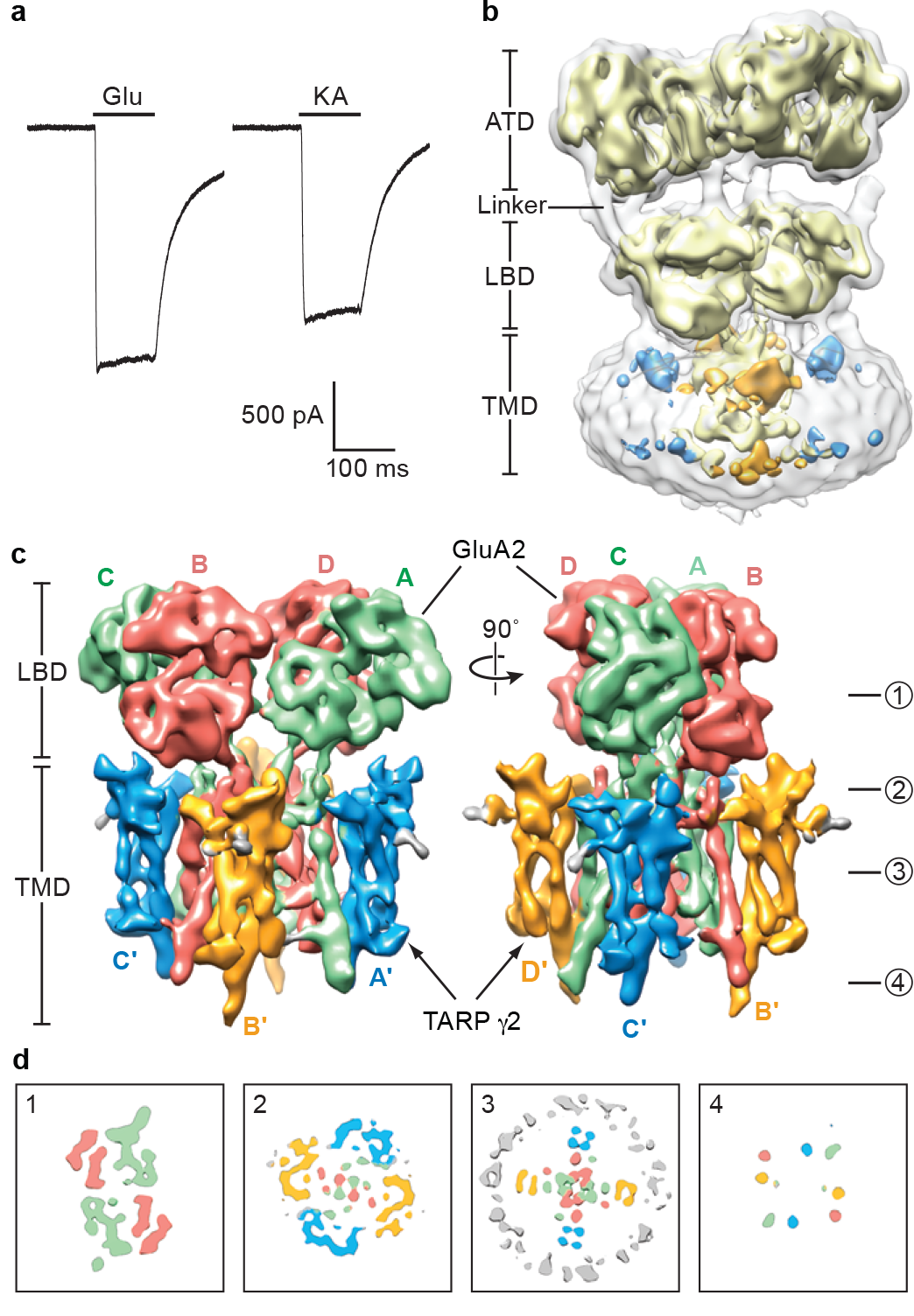
Function and reconstruction of GluA2-TARP γ2 complex. a, Whole-cell patch clamp recordings from cells expressing the GluA2 – TARP γ2 complex. A representative pair of currents recorded using the same cell is shown. The ratio between steady-state currents evoked by kainate and glutamate is 0.80±0.02 (mean ± standard deviation, n=5). b, Initial 3D-reconstruction of GluA2-TARP γ2 complex contoured at lower (outer) and higher (inner) threshold levels showing distinct features for the receptor and associated TARP. c, Refined 3D-reconstruction focused on LBD and TMD layers, where the ATDs were excluded from refinement. A/C and B/D subunits of the GluA2 receptor are in green and red, respectively; TARP γ2 associated with receptor A/C and B/D subunits are in blue and gold respectively. d, Cross-sections of the EM map at LBD layer, TARP-LBD interface layer, TMD layer and C-terminal layer at indicated ‘height’, with density features colored as in panel (c).

Three-dimensional reconstruction of the receptor-TARP complex without the imposition of symmetry revealed an overall architecture consistent with previous crystal and cryo-EM structures of the antagonist-bound GluA2 receptor^10,13^ (Fig. 1b). The initial 3D classification yielded four classes, one of which had four protrusions on the extracellular side of the detergent micelle, related by an approximate 4-fold axis of symmetry, and was composed of the largest number of particles. The remaining 3 classes had poorly resolved features associated with the extracellular domains and did not exhibit 4-fold symmetric protrusions from the micelle, features associated with the presence of TARP subunits, and thus were excluded from the analysis (Extended Data Fig. 2). Further studies, and larger data sets, will be required to elucidate the structures of additional structural classes of the receptor – TARP complex.

To improve the density of the TARPs and the structural features of receptor-TARP interactions, we carried out focused refinement of the LBD and TMD layers^28^, masking the conformationally heterogeneous ATD layer, with application of C2 symmetry coincident with the two-fold axis that relates the LBD dimers and the four-fold axis of the TMD, in the subsequent 3D reconstructions and refinements (Extended Data Fig. 2). The resulting density map has an estimated resolution of 7.3 Å (Extended Data Fig. 3) and illustrates hallmark features of the LBD clamshells and the receptor TMD. Most importantly, the density map clearly reveals the presence of four TARPs, arranged with four-fold symmetry, surrounding the exterior of the receptor TMD, consistent with a fully saturated receptor-TARP complex (Fig. 1c). The receptor-TARP complex features a similar symmetry mismatch between the two-fold related LBD layer and four-fold related TMD layer as found in isolated receptor (Fig. 1d), with TARP subunit pairs A’/C’ and B’/D’ ‘underneath’ the A/C and B/D LBDs, respectively. We further note that density for the full-length receptor M4 and carboxy-terminal TARP TM helices extend into the cytoplasm (Fig. 1d).

To generate the structural model of GluA2 receptor – TARP γ2 complex, we extracted individual LBDs and the intact TMD from the MPQX-bound GluA2 crystal structure^10^ and fit them into the cryo-EM density as rigid bodies (Fig. 2a and 2b). Manual adjustments of secondary structure elements were applied where there was supporting density, followed by fitting of the preM1 and M3-S2 linkers (Fig. 2c and 2d). The S2-M4 linkers were not visible in the density maps. The cryo-EM density for the ATDs was poorly resolved, in line with their flexibility. Thus, we did not focus on optimizing the density of the ATD layer, concentrating instead on the crucial LBD, TMD and TARP regions. The degree of LBD clam-shell opening is similar to the MPQX-bound full-length receptor structure^10^, confirming that the complex is stabilized in an antagonist-bound state (Fig. 2a).

**Figure 2.**
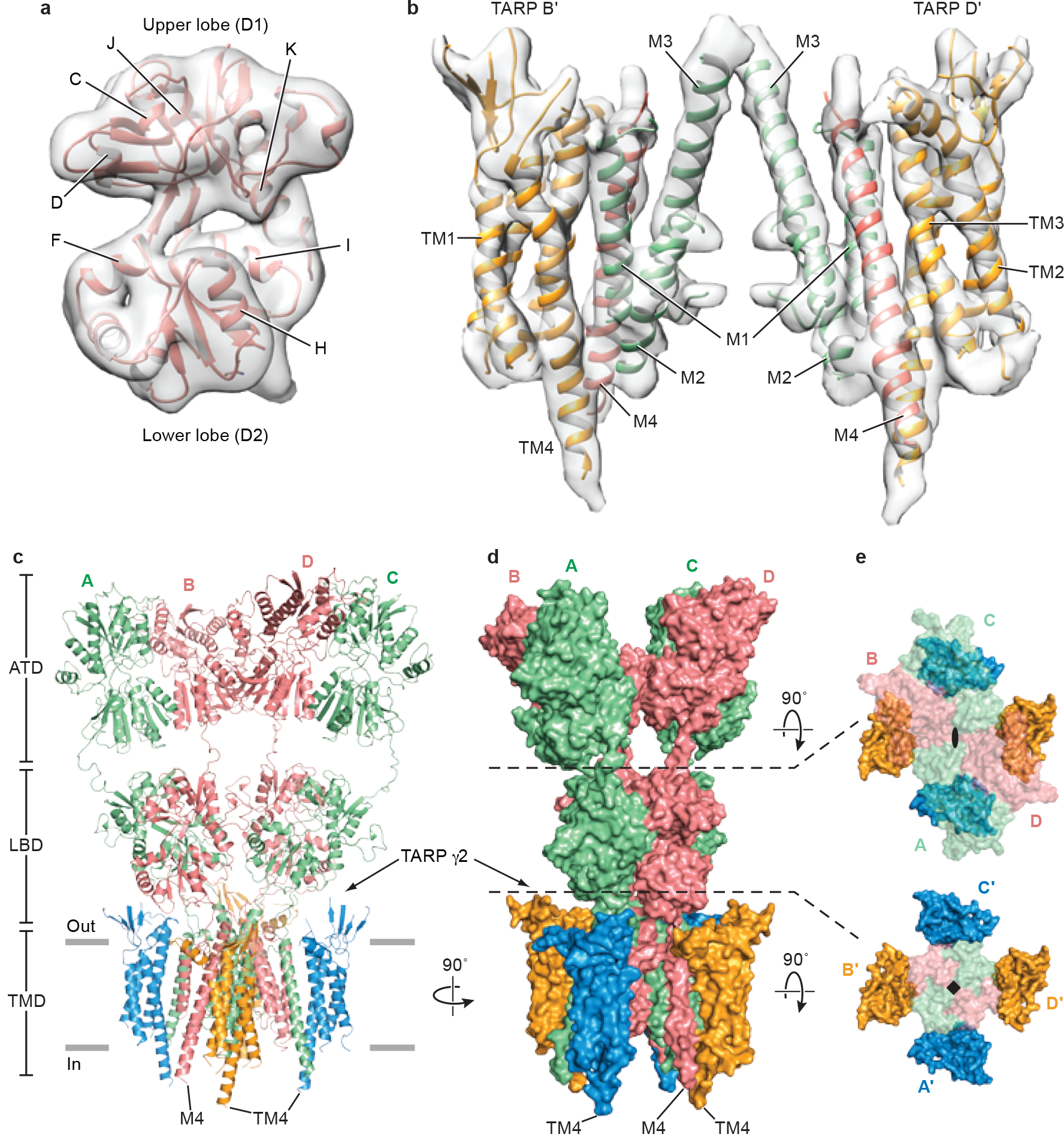
Structure of GluA2 receptor in complex with TARP γ2. Cryo-EM density maps for LBD and TM regions of the complex in panels **a** and **b** respectively. A’/C’ TARPs and associated receptor TMs were omitted for clarity. Ribbon diagram and surface representation of the complex in panels **c** and **d**. The carboxy-termini of selected TARP helices (TM4) and receptor helices (M4) are labeled. **e**, Two “top-down views”, in which elements of the structure above the indicated dashed lines have been omitted, for clarity, from the figures. The GluA2 receptor is shown in transparent surface representation to allow visualization of the TARP subunits.

Because *de novo* structure determination for TARP was not feasible at the resolution of this study, we generated a homology model of the TARP γ2 using the claudin-19 crystal structure as a template^4^ (Extended Data Fig. 4a). Rigid body fitting of the TARP model in the density map was unambiguous, driven by strong helical density for the TARP TMs and consistent with computational analysis of the TARP density by the program SSEhunter^29^ (Extended Data Fig. 4b-d). The particularly long TM4 of TARP, which protrudes into the cytoplasm, provides an additional structural landmark by which to validate the fitting of the TARP model to the experimental density map. Whereas there was strong density to support the presence of a TARP β-sheet on the extracellular side of the membrane, like that found in claudin-19^4^, little density was found for the ‘loop’ connecting the β1 and β2 strand, or that between TM3 and TM4 (Extend Data Fig. 4a). These loops were therefore excluded from the homology model (Extended Data Fig. 4c and 4d). In addition, a short helix (α1) was placed into the tube-like density adjacent to TM2, an assignment that was supported by the SSEhunter scores^29^ and sequence based secondary structural prediction^30^ (Extended Data Fig. 4b). The final TARP model resembles a forearm and partially open right hand, with the TMD representing the arm, the β-sheet representing the palm, and the short α1 helix representing the thumb (Extended Data Fig. 4b and 4c).

The closer proximity between LBDs and TARPs at the B/D positions compared to the A/C positions suggest that the A’/C’ and B’/D’ TARPs play non-equivalent roles in the modulation of receptor activities (Figure 2c and 2d). Indeed, the lower lobe of the B/D LBDs has been proposed to play a greater role in ion channel gating^31^. Nevertheless, it is possible that movements in the LBD layer upon agonist binding could cause the LBDs to engage the A’/C’ TARPs, a hypothesis supported by evidence that binding of 4 TARPs leads to greater activation by the partial agonist kainate^24^, highlighting the functional significance of TARP subunits in all four positions.

The interactions between TARP and receptor are comprised of two components:TM-TM interactions and TARP extracellular domain (ECD)-receptor LBD interactions. The TM-TM interactions at both A/C and B/D positions are equivalent, obeying the four-fold symmetry of the TMD (Fig. 2e). The TM3 and TM4 of TARP form extensive hydrophobic interactions with M1 and M2 from one GluA2 subunit and with M4 from the adjacent subunit (Fig. 3a and Extended Data Fig. 6), thus suggesting one structural mechanism by which TARPs can modulate the properties of the ion channel. Given the nearly identical receptor TMD structure observed in apo and pre-open states, the TARP TMDs likely remain bound to receptor in a similar fashion in activated states. By contrast, there are no direct contacts between the TARP thumb and palm and the receptor LBDs (Fig. 3b), although visualization of such interactions may be limited by the resolution of the reconstructions as well as inherent TARP flexibility. Nonetheless, a conserved acidic region spanning residues 85-95 (sequence:EDADYEADTAE) is present in the TARP extracellular ‘loops’ adjacent to the α1 helix (Extended Data Fig. 4a), poised to interact with several positively charged residues on the lower lobe of the LBD, including the “KGK” sequence at residues 718-720 of the receptor^32^ (Fig. 3b). While these elements of structure may be too distant to form salt bridges in this antagonist-bound state, we speculate such interactions could take place in pre-open or activated states (Extended Data Fig. 7), consistent with the importance of both TARP ECD and the LBD “KGK” motif, as well as a ‘lowering’ of the receptor LBD toward the membrane upon receptor activation^11,12^.

Elucidation of the architecture of the GluA2 receptor-TARP γ2 complex was facilitated by FSEC-based screening, which shows that digitonin stabilizes the receptor-TARP complex. Analysis of the structure by single particle cryo-EM illuminates how four TARPs encircle the receptor TMD and participate in extensive interactions with receptor TM helices, thus demonstrating the importance of non-polar contacts in complex formation. The acidic, partially open TARP ‘palms’ are positioned underneath basic motifs on the lower lobes of the LBDs (Fig. 4a), illustrating how complementary electrostatic interactions also contribute to receptor-TARP interactions (Fig. 4b). By juxtaposition of the TARP ‘palms’ underneath the LBD ‘clamshells’, TARPs are ideally positioned to modulate domain closure and thus efficacy of partial agonists.Moreover, the A’/C’ TARP proximity to the LBD dimer interface and the closeness of the B’/D’ pair to the LBD dimer-dimer interface suggest how TARPs might modulate the activity of positive allosteric modulators and the modal gating properties of the receptor, respectively (Fig. 4a and 4b). We further speculate that the spatially distinct pairs of TARPs offer a structural explanation for biexponential kinetics deactivation and desensitization of the receptor-TARP complex. Lastly, TARP TMD extensively interacts with receptor TMD including the pore helix M2, stabilizing the M2 helix and selectivity filter, thereby suggesting a mechanism for TARP-modulation of receptor pore properties.

**Figure 3.**
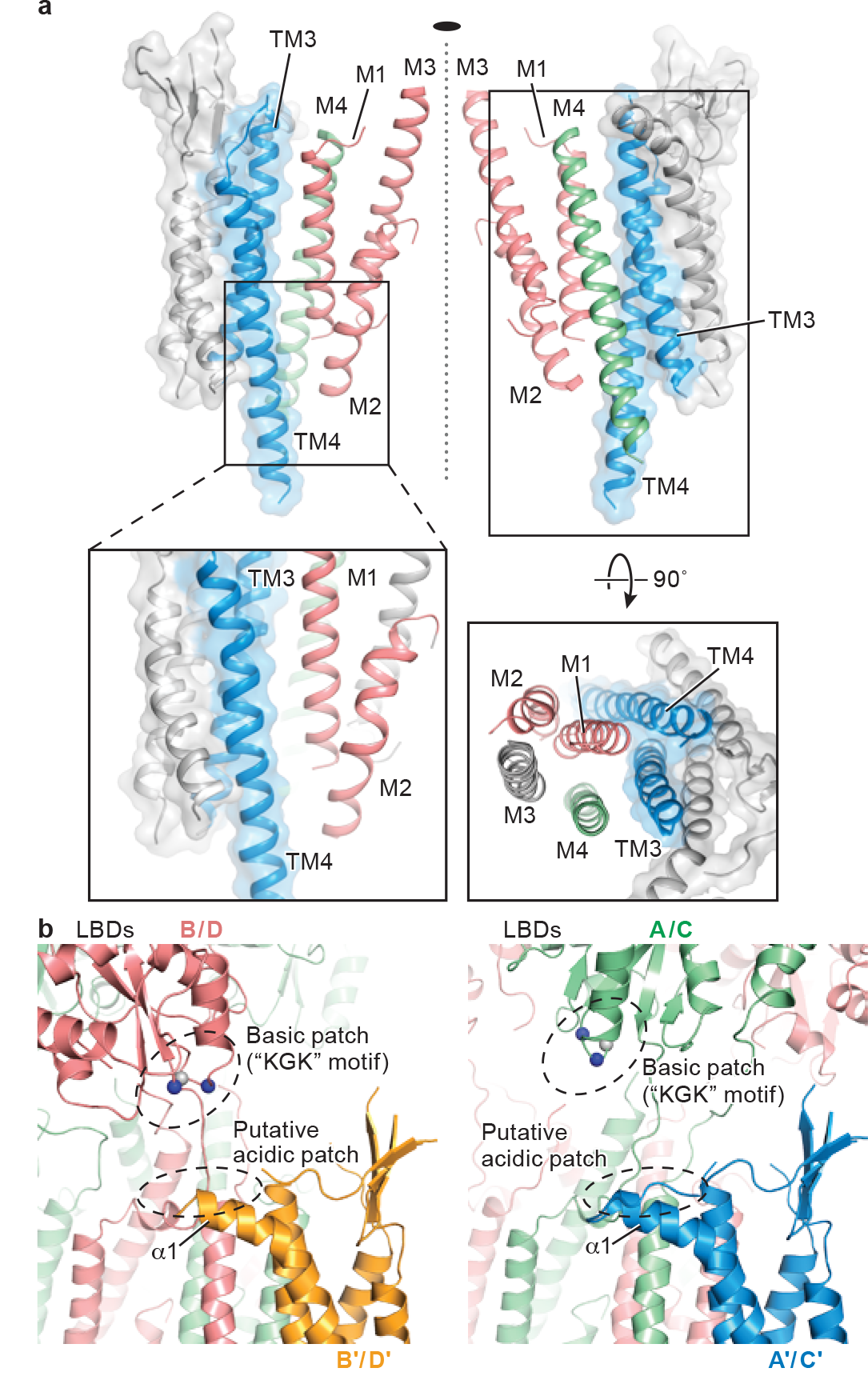
Interactions between GluA2 receptor and TARP γ2. **a**, TM-TM interactions between TARP γ2 and receptor at A/C positions. TARP γ2 is shown in ribbon diagram in transparent surface representation, while helices participating in interactions with receptor are highlighted in color. The central axis of the ion channel pore is indicated by a dashed line. A close-up view emphasizes hydrophobic interactions between TARP and receptor, whereas a top-down view illustrates that all TM helices but M3 of the receptor interact with TARP. TM helices from selected receptor subunits were omitted for clarity. **b**, Interactions between TARP γ2 ECD and receptor LBD at the A/C positions differ from interactions at the B/D positions. TARP γ2 ECD and receptor LBD are in closer proximity in the B/D positions (right) than in the A/C positions (left). Approximate locations where positively charged residue cluster from the receptor and negatively charged residue cluster from TARP are indicated by dashed circles. The Cα atoms of the “KGK” motif (718-720) are shown as spheres. Lysine and glycine Cαs are colored in blue and grey respectively.

**Figure 4.**
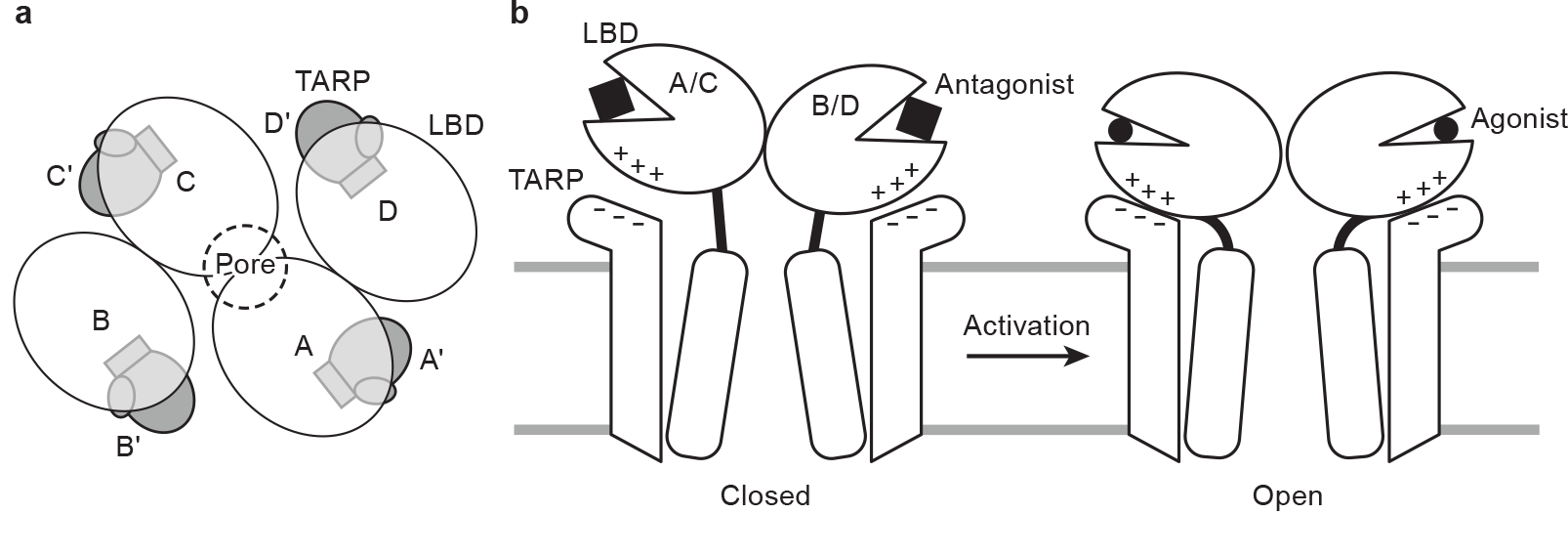
Mechanism for TARP γ2 modulation of receptor gating. **a**, TARP γ2 subunits resemble partially opened palms and are positioned “underneath” receptor LBDs in the antagonist-bound state. **b**, During receptor activation, TARP “palms” engage with receptor LBD to stabilize intra-dimer and inter-dimer interfaces, modulating receptor activation, deactivation and desensitization. An extracellular loop of TARP γ2 rich in negatively charged residues facilitates the motion of receptor LBD lower lobe rich in positively charged residues upon receptor activation, whereas TARP γ2 TMD directly interacts with receptor TMD including the pore-lining M2 helices, leading to modulation of receptor pore properties.

## Materials and methods

### Electrophysiology

Electrophysiology experiments were performed using a stable cell line (Clone #10) that constitutively expresses full-length wild-type TARP γ2 and the C-terminally FLAG-tagged GluA2 AMPA receptor (flop isoform, arginine for the Q/R site) under control of the TerON promoter^26^. Whole-cell recordings were carried out 10-18 hours after induction of GluA2 receptor expression with 7.5 μg/ml doxycycline. Pipettes were pulled and polished to 2-3 MO resistance and filled with internal solution containing 75 mM CsCl, 75 mM CsF, 5 mM EGTA and 10 mM HEPES pH 7.3. External solution contained 160 mM NaCl, 2.4 mM KCl, 4 mM CaCl_2_, 4 mM MgCl_2_, 10 mM HEPES pH 7.3 and 10 μM (R,R)-2b, a positive modulator that blocks desensitization^33^. To allow efficient binding of (R,R)-2b, each cell was perfused in external solution for one minute before currents were elicited by 3 mM of L-glutamate and 0.6 mM kainate, individually, with a one-minute wash step in between. Ratios of glutamate and kainate-evoked currents determined in five independent experiments were subjected to statistical analysis using Origin.

### Expression and purification

The AMPA receptor-TARP γ2 complex was expressed using clone #10 cells adapted to grow in suspension^26^, using freestyle 293 expression medium (gibco, life technologies) supplemented with 2% (v/v) fetal bovine serum (gibco, life technologies) and selection antibiotics (125 μg/ml zeocin, 150 μg/ml hygromycin, and 125 μg/ml neomycin). GluA2 (flop variant, arginine at the Q/R site) expression was induced by addition of 7.5 μg/ml doxycycline at a cell density of 2×10^6^ cells/ml. Subsequently, MPQX^27^ was added to the media (final concentration of 200 nM) to prevent cytotoxicity due to receptor overexpression. Cells were harvested by centrifugation 30~35 hours after induction and homogenized by sonication. After removal of cell debris by centrifugation at 1,200 × g (15 min at 4 °), the supernatant was subjected to ultracentrifugation at 100,000 × g for 1 hour to collect the membrane fraction.

The membrane fraction was resuspended and solubilized in TBS buffer (20 mM Tris, pH 8.0, 150 mM NaCl) containing 1% (w/v) digitonin (Calbiochem) and 1 μM MPQX for 2 hours at 4 °. Insoluble material was removed by ultracentrifugation at 100,000 x g for 1 hour, and the supernatant was passed through an anti-FLAG immunoaffinity column pre-equilibrated with buffer P (20 mM Tris pH 8.0, 150 mM NaCl, 0.1% (w/v) digitonin, 1 μM MPQX), followed by a wash step using 10 column volumes of buffer P. The FLAG-tagged GluA2 receptor in complex with TARP was eluted with buffer P supplemented with 0.5 mg/ml FLAG peptide. The eluted complex was concentrated and further purified by size-exclusion chromatography (SEC) using a Superose 6 10/300 GL column (GE Healthcare) equilibrated in Buffer P. Peak fractions were pooled and concentrated to 3 mg/ml using Amicon 100-kDa cutoff concentrator (EMD Millipore) for subsequent biochemical analysis and cryo-EM studies.

### Cryo-EM data acquisition

A droplet of 2.5 μl of purified AMPAR-TARP γ2 complex at 3 mg/ml was placed on quantifoil 0.6-1.0 Au 200 mesh grids glow discharged at 30 mA for 120s. The grid was then blotted for 2~3 s at 22 °C under conditions of 100% humidity, and flash-frozen in liquid ethane.

Cryo-EM data were collected on a 300 kV Titan Krios microscope (FEI) using a K2 camera (Gatan) positioned post a GIF quantum energy filter (Gatan). The energy filter was set to a 20 eV slit and a 70 μm objective aperture was used. Micrographs were recorded in superresolution mode at a magnified physical pixel size of 1.35 Å, with the defocus ranging from -1.5 to -2.5 μm. Recorded at a dose rate of 8.3 e^-^/pix/s, each micrograph consisted of 40 dose-fractionated frames. Each frame was exposed for 0.3 s, resulting in a total exposure time of 12 s and total dose of 55 e^-^/Å^2^.

### Image processing

A total of 2675 micrographs were subjected to motion correction with Unblur^34^. The CTF parameters for each micrograph were determined by CTFFIND3^35^ and particles were picked using DoG picker^36^. Several rounds of 2D classification were used to remove ice contamination, micelles, disassociated or disordered protein and other false positives. The large number of particles discarded was likely a consequence of using DoGPicker with a fairly large threshold range; earlier attempts using template-based correlation manifested in an orientation bias during 2D classification. In this way, 2D classification also served as an opportunity to assess how well the selected particle orientations were distributed (Extended Data Fig. 3a). Rounds of 2D classification were repeated until the remaining classes had features recognizable from a comparison with an ensemble of 2D projections calculated by using the crystal structure of the antagonist-bound receptor. From an initial set of 257,378 putative particles, 61,539 particles were selected for subsequent 3D classification (Extended Data Fig. 1).

Initially, this subset of particles was classified into four classes using a reference model generated from the GluA2 X-ray structure of the MPQX-bound state^10^ (PDB code:3KG2), which had been low-pass filtered to 60 Å. The most populated 3D class, containing 49% of total particles, featured four “bumps” on the extracellular side of the detergent micelle, and was subjected to further 3D refinement using a soft mask focused on the LBD and TMD domains, with C2 symmetry imposed^37^. This further improved the quality of the density map, allowing the final map to reach 7.3 Å resolution as estimated by Fourier shell correlation between two independently refined half maps. Failing to show any putative TARP features and having poorly resolved features for the extracellular domains, the remaining three 3D classes possibly represented receptor free of TARP or even residual false positives, and were therefore excluded from subsequent analysis. All 2D and 3D classifications and refinements above were performed in RELION 1.4^38^.

### Structural modeling

The structural modeling for the GluA2-TARP γ2 complex was comprised of rigid-body fitting of LBDs and TMDs extracted from the MPQX-bound GluA2 crystal structure (PDB code:3KG2) into the cryo-EM density, followed by fitting of a homologous TARP model generated by SWISS-MODEL^39^ using the crystal structure of claudin-19^4^ (PDB code:3X29) and sequence alignment performed with Clustal Omega^40^. A 25.7% sequence similarity (11.3% identity) between claudin-19 and TARP γ2 was determined by Sequence Manipulation Suite^41^. Docked as a rigid body, the derived TARP homology model was refined in real space against the density map using COOT^42^ guided by the resolved helical density of the TARP TMs, and the density consistent with the conserved β-sheet on the extracellular side of the micelle. Furthermore, the TARP TM4 helical density was observed to protrude from the cytoplasmic side of the micelle, further assisting in the fitting of the TARP helices to the density map. The entire model was then improved by manual adjustments including removing several loop regions outside of density, local rigid-body fitting of individual helices into density, extension of the TARP TM4 helix by 14-residues and positioning of a short helix (α1) adjacent to TM2 supported by secondary structure prediction (Jpred4^30^). The extension of the TARP TM4 helix was justified by strong density in the experimental density maps consistent with continuation of the α-helix, scoring in SSEhunter consistent with an α-helix, and prediction of these residues in an α-helical conformation by secondary structure prediction (Jpred4^30^).

To validate the fitting and the placement of TARP TM4 extension and α1 helix, we used SSEhunter^29^ to verify the secondary structure assignment against the EM density. To do this, the putative TARP density was extracted in Chimera^43^ using Segger^44^. SSEhunter analysis resulted in a series of pseudoatoms located on the skeleton of the density map, each assigned with a score. The positive scores at TM and α1 helix region and negative scores at the β-sheet region confirmed the secondary structure elements present in the TARP model.

The final map was put into a large P1 unit cell (*a* = *b* = *c* = 405 Å α = β = γ = 90° and structure factors were calculated in PHENIX^45^. The complex model of GluA2 receptor (LBD and TMD, residues 400-551, 570-593, 596-782, 787-832) – TARP (residues 6-38, 56-68, 72-82, (AC:84-126)/(BD:91-125), 131-162, 174-215; see also Extended Data Fig. 4) was then refined against structure factors derived from the density map using phenix.real_space_refine^45^. Secondary structure, 2-fold NCS and Ramachandran restraints were applied throughout the entire refinement. After refinement, map CC between model and EM map was 0.716, indicative of a reasonable fit at the present resolution. The resulting model was also used to calculate a model-map FSC curve, which agreed well with the gold-standard FSCs generated during the RELION refinement (Extended Data Figure 3c). The final model has good stereochemistry, as evaluated using MolProbity (Extended Data Table 1).

All of the figures were prepared with Pymol^46^, UCSF Chimera^43^ and Prism 5.

## ACKNOWLEDGEMENTS

We thank T. Nakagawa for providing the clone #10 cell line Z. H. Yu R. Huang and C. Hong (Janelia Campus) for assistance with microscope operation and data collection and the Advanced Computing Center (OHSU) for computational support. We are grateful to the Multiscale Microscopy Core (OHSU) for support with microscopy, and L. Vaskalis for assistance with figures H. Owen for help with proofreading and other Gouaux laboratory members for helpful discussions. S.C. is supported by an American Heart Association postdoctoral fellowship (16POST27790099). This work was supported by the NIH (E.G., NS038631). E.G. is an investigator with the Howard Hughes Medical Institute.

### AUTHOR CONTRIBUTIONS

Y.Z., S.C. and E.G designed the project. Y.Z. and S.C. performed sample preparation and cryo-EM data collection. Y.Z. and C.Y. analyzed the data. I.B. performed electrophysiology experiments. Y.Z., S.C., C.Y. and E.G. wrote the manuscript with input from I.B.

### AUTHOR INFORMATION

The three-dimensional cryo-EM density map and the coordinate for the structure of GluA2 AMPA receptor – TARP γ2 have been deposited in the EM Database and Protein Data Bank under the accession codes EMD-8256 and 5KK2, respectively. The authors declare no competing financial interests. Correspondence and requests for material should be addressed to E.G.

## EXTENDED DATA FIGURE LEGENDS

**Extended Data Figure 1. Digitonin is a suitable detergent for the purification of GluA2 receptor – TARP γ2 complex. a**, The receptor-TARP complex in digitonin disassociates when diluted into DDM. A complex composed of the GluA2 receptor and GFP-tagged TARP γ2 was diluted in digitonin (green) or DDM (red) before being subjected to GFP-tuned FSEC analysis. **b**, Coomassie blue-stained SDS-PAGE gel of the purified complex. **c**, Tryptophan-tuned FSEC profile of the purified complex was comprised of a major peak containing the tetrameric complex and a minor shoulder, the latter suggestive of either incompletely assembled or partially dissociated complexes. Only the full-size, tetrameric species was used for single particle cryo-EM analysis. **d**, A representative, motion-corrected micrograph of the GluA2 receptor – TARP γ2 complex is shown. A few distinct complexes with the characteristic capital Y shape of the non-desensitized state of the AMPA receptor are circled. **e**, Representative 2D-class averages showing a range of projections of the receptor – TARP γ2 complex.

**Extended Data Figure 2. The work-flow of cryo-EM data processing**. The raw dataset used in this study was composed of 2,675 micrographs. Particles (257,378) were picked from motion-corrected and CTF-estimated micrographs for subsequent classifications. After multiple rounds of 2D classification, the remaining 61,539 particles were subjected to several rounds of 3D classification. Initial 3D classification yielded four major classes, where the most populated one contained 49% of total particles. An initial 3D reconstruction without imposed symmetry resulted in a moderate resolution at 9.6 Å. With C2 symmetry imposed, subsequent 3D refinement focused on the LBD and TMD layer improved the density map, allowing a reconstruction at 7.6 Å resolution. An additional two iterations of 3D classification and refinement using updated map as the reference further improved the reconstruction. The resolution of the final cryo-EM density map was estimated to be 7.3 Å.

**Extended Data Figure 3. Statistics for the cryo-EM reconstruction. a**, Euler angular distribution of all particles included in the final 3D reconstruction. The number of particles viewed from each specific orientation was indicated by the size of the corresponding sphere. **b**, Gold-standard FSC curves calculated between two independently refined half-maps before (red) and after (blue) post-processing, overlaid with FSC curve calculated between cryo-EM density map and structural model.

**Extended Data Figure 4. Structures of the TARP γ2 subunits in the context of the respective cryo-EM density map. a**, Sequence alignment between TARP 72 and claudin-19 calculated using Clustal omega. Also shown above the alignments are the secondary structure elements of TARP γ2 based on the model reported here, and below the aligned sequences are the secondary structure elements derived from the claudin-19 crystal structure. The ECD region rich in negative charges is conserved throughout the TARP family and highlighted in red. **b**, EM density for B’ TARP and pseudo-atoms placed by SSEhunter, each colored according to a calculated secondary structure score. Positive and negative scores indicate α-helix and β-sheet propensity respectively. Dashed-line circles a map region where high scores were found, suggesting the presence of helical structure. A scale bar ranging from a maximum positive value (α-helix) to the minimum negative score (β-strand) is shown. **c**, The A’ TARP of the A’/C’ pair. **d**, The B’ TARP of the B’/D’ pair. The first and last visible residues Arg6 and Thr215 were labeled. Secondary structure elements were color-coded as in panel a.

**Extended Data Figure 5. Structural comparison between TARP γ2 and claudin-19**. A superposition of the TARP γ2 structure (in blue) derived from this study and claudin-19 (in grey) is consistent in the conserved overall fold, with the exception that there is a short α1 helix present only in TARP γ2.

**Extended Data Figure 6. Cryo-EM density map for the pore-helix of the GluA2 receptor**.Clear density (blue mesh) is present for the pore-lining M2 helices, secondary structure elements that are weak or absent in all previous crystal structures. The N-terminus of each pore helix is involved in extensive interactions with TM4 from TARP subunits and we suggest that interactions of receptor TM helices that include M2, with TARP TM helices, stabilize the ion channel pore.

**Extended Data Figure 7. Possible interaction between receptor LBD and TARP in an active state**. Shown on the left is a “top-down view” of MPQX-bound receptor-TARP complex structure (in color) superimposed with the crystal structure of an active state GluA2 receptor (in grey) in complex with a partial agonist, fluorowilliardiine (FW) and a positive allosteric modulator, (R, R)-2b, using the central M3 helices as a reference. ATDs and LBDs were omitted for clarity. The modeled pseudo-complex consisting of TARPs and the active state receptor illustrates a possible mechanism for how TARP interacts with receptor LBD during activation. Enlarged views of the MPQX-bound complex structure and FW/(R, R)-2b bound complex model were shown side by side at both D’ and A’ TARP positions. LBD helices and a S2-M4 linker were labeled according to convention.

## EXTENDED DATA TABLE TITLES AND LEGENDS

**Extended Data Table 1.** Statistics of cryo-EM data collection, 3D reconstruction and model refinement.

|  |  |
| --- | --- |
| **Data collection/processing** |  |
| Microscope | Krios |
| Voltage (kV) | 300 |
| Camera | Gatan K2 |
| Camera mode | Super-resolution |
| Defocus range ($\mu\text{m}$) | -1.5 ~ -2.5 |
| Exposure time (s) | 12 |
| Dose rate ($e^-/\text{pixel/s}$) | 8.3 |
| Magnified Pixel size ($\text{\AA}$) | 1.35 |
| **Reconstruction** |  |
| Software | RELION 1.4 |
| Symmetry | C2 |
| Particles refined | 26,297 |
| Resolution (unmasked, $\text{\AA}$) | 9.0 |
| Resolution (masked, $\text{\AA}$) | 7.3 |
| Map sharpening B-factor ($\text{\AA}^2$) | -600 |
| **Model Statistics** |  |
| Protein residues | 2309 |
| Map CC | 0.716 |
| Resolution (FSC=0.5, $\text{\AA}$) | 8.0 |
| MolProbity score | 2.49 |
| Cβ deviations | 0 |
| Ramachandran |  |
| Outliers | 0.27% |
| Favored | 88.28% |
| RMS deviations |  |
| Bond length | 0.008 |
| Bond angles | 1.26 |

